# Heterochronic Uncoupling of Brain and Face Morphogenesis Drives Premature Fusion in Ciliary Transition Zone Mutants

**DOI:** 10.64898/2026.10.19.760125

**Authors:** Diane Hu, Jay Devine, Wei Liu, Ralph S. Marcucio, Benedikt Hallgrímsson, Rebecca M. Green, Nathan M. Young

## Abstract

Normal facial development is highly coordinated with embryonic brain expansion, suggesting the forebrain acts as a physical scaffold dictating the spatial timing of facial prominence apposition and fusion. To test this idea, we compared *B9d1* ciliary transition zone mutants, which exhibit altered Hedgehog signaling and premature facial fusion, to a normal murine developmental morphospace. At e11.5, mutant brains are significantly reduced in size, yet intrinsic facial prominence size remains largely normal. Projecting e11.5 and e12.5 mutants into the normal morphospace reveals a severe heterochronic uncoupling: the delayed, narrowed neurocranium mechanically forces normal-sized facial prominences into premature convergence. These results indicate that when brain growth slows, facial prominences initiate contact earlier in restricted spatial bounds. Conversely, this model predicts that an accelerated, expanded brain may act as a physical wedge, increasing the likelihood of cleft lip. By mathematically modeling these structural dependencies, we propose that palatogenesis operates on a continuous phenotypic spectrum, where premature fusion and cleft lip represent alternative physical outcomes conditioned on the underlying brain developmental trajectory.

**Summary Statement:** 3D developmental morphospace modeling reveals that delayed brain growth in transition zone mutants imposes a mechanical constraint, physically forcing normal-sized facial prominences into premature midline fusion.

## Introduction

Normal craniofacial development requires the precise spatiotemporal choreography of multiple tissues to produce the highly integrated structures of the vertebrate head (Chai & Maxson, 2006). In particular, the formation of the primary palate and upper jaw depends on the outgrowth, contact, and subsequent fusion of the paired maxillary, medial and lateral nasal facial prominences. Given the complex spatiotemporal orchestration involved in palatal development, even minor deviations in facial prominence growth or positioning frequently propagate into severe downstream dysmorphologies (Dixon *et al*., 2011). Unsurprisingly then, cleft lip is one of the most common congenital anomalies in humans, representing a failure of the initially independent facial prominences to successfully integrate into the upper jaw (Jiang *et al*., 2006).

Historically, efforts to understand the etiology of cleft lip have focused heavily on factors intrinsic to the prominences, linking fusion failure to localized reductions in proliferation, cell survival, or absolute prominence size (Trasler, 1968; Diewert *et al*., 1993; Wang *et al*., 1995; Diewert & Lozanoff, 2002; Young *et al*., 2007). For example, in the *Wnt1-Cre; Srsf3 fl/fl* mutant model, widespread neural crest cell death creates an intrinsic volume deficiency that directly causes a severe midfacial cleft (Dennison *et al*., 2021). In contrast, the recently recovered *Unicorn* mouse model reveals that identical severe clefting can arise from a pure geometric vector mismatch (a laterally shifted frontonasal ectodermal zone) despite absolutely normal cellular proliferation and viability (Lantz *et al*., 2020). Because pure geometric mismatches (*Unicorn*) and classic volume deficiencies (*Srsf3*) yield similar clefting phenotypes, the field requires a system-wide structural model to map both macro-level and cellular constraints.

While cellular dynamics localized to the facial prominence are undoubtedly crucial to normal palatal development, explicit comparative modeling of facial growth across amniotes reveals an equally critical, deeply constrained, system-wide dynamic. Using three-dimensional geometric morphometrics (3DGM) to construct a continuous developmental morphospace, we previously demonstrated that despite early and late macroevolutionary diversity, amniote facial shape trajectories predictably converge at primary palatal fusion before diverging into adult species-specific proportions (Young *et al*., 2014). Building upon this comparative model, our recent continuous sliding-window morphometric analysis demonstrated that embryonic craniofacial development in the chicken (*Gallus gallus*) is characterized by both a tightly integrated biphasic trajectory and a collapse in spatial variance coincident with palatogenesis (Hu *et al*., 2026). This structural bottleneck indicates that successful facial fusion requires embryos navigate a constrained “shape window.”

Importantly, facial prominences do not develop in isolation. Their growth is dependent on the broader structural environment, and most notably, the expanding forebrain. Our previous work in the chicken indicates that brain shape executes a similar highly synchronized trajectory reversal coincident with facial fusion (Hu *et al*., 2026). This coordination suggests that the embryonic brain serves a dual role in facial morphogenesis: it acts both as a critical molecular signaling center and as a physical, expanding scaffold (DeMyer *et al*., 1964; Muenke *et al*., 1994; Boughner *et al*., 2009; Marcucio *et al*., 2011; Chong *et al*., 2012; Hu *et al*., 2015). As the forebrain increases in volume, the expansion of neuroepithelial surface area alters the physical distance that the growing facial prominences must cross to achieve contact and fusion. Consequently, intrinsic brain development likely serves as a critical spatiotemporal dependency for successful palatogenesis.

Based on this “phenotypic landscape” model of brain-face integration, we hypothesize that fusion events exist along a phenotypic spectrum conditioned on the trajectory of brain development. In normal brain development, facial contact and fusion is dependent on factors internal to the prominences themselves. However, if the brain develops faster than normal, the time in which facial prominences can contact will be reduced, separating the midface before fusion occurs and increasing the likelihood of cleft lip. Conversely, if brain development is delayed or slowed relative to the face, the facial prominences will initiate contact earlier and in abnormal spatial locations, leading to premature apposition and potentially fusion.

Testing the predictions of this “phenotypic landscape” model requires disrupting the integrated signaling and growth relationship between the brain and the face. The primary cilium is an essential organelle for Hedgehog (SHH) signal transduction, functioning as the localized nexus where the receptor Patched restricts Smoothened (Smo) accumulation, thereby regulating the proteolytic processing of Gli transcription factors into their activator or repressor forms (Rohatgi *et al*., 2007; Bangs & Anderson, 2017). A key component of the primary cilia is the transition zone complex, which includes proteins such as B9d1, Tctn2, Tmem231, and Mks6. Mutations to the transition zone complex disrupt ciliary gating, altering SHH signaling, and ultimately leading to a spectrum of severe craniofacial and neural tube defects with overlapping effects (Dowdle *et al*., 2011; Schock & Brugmann, 2017; Abrams & Reiter, 2021).

To mechanistically test the role of the forebrain in dictating spatial thresholds for palatal fusion, we utilized the B9d1 mutant mouse model, which exhibits characteristic alterations in SHH signaling and profound craniofacial abnormalities, including premature apposition and fusion of the facial prominences in the midline. By mapping high-resolution micro-computed tomography (µCT) data of B9d1 mutants onto a highly detailed, continuous developmental morphospace of normal mouse craniofacial morphogenesis, we sought to determine how localized transition zone mutations disrupt integrated trajectories of the brain and face. Our analyses demonstrate that B9d1 mutants exhibit heterochronic uncoupling, driven primarily by altered brain shape and size, which collapses the structural landscape and mechanistically forces the premature apposition of normal-sized facial prominences. These results provide a quantitative framework for understanding how brain developmental trajectories conditionally dictate the mechanical thresholds for both normal palatogenesis and structural pathology.

## Results

### Phenotypic Convergence of Ciliary Transition Zone Mutants at the Fusion Window

To determine whether disruption of the transition zone complex yields a generalized loss of structural integrity or a rigidly canalized developmental trajectory, we first evaluated the unconstrained shape variance of external craniofacial morphology across a multi-mutant cohort at embryonic day 11.5 (e11.5). Principal Components Analysis (PCA) was performed on external facial and whole-head landmarks of null mutants (*B9d1^-/-^*), wild-type (*B9d1^+/+^*), and heterozygous (*B9d1^+/-^*) littermates, alongside three additional ciliary transition zone null mutant models (*Tmem231^-/-^*, *Mks6^-/-^*, and *Tctn2^-/-^*). The resulting morphospace revealed a distinct phenotypic convergence [**Figure 1A**]. Wildtype and heterozygous individuals clustered together, establishing the boundaries of normal morphological variation. In contrast, all four transition zone null mutant models were separated into a distinct region of morphospace characterized by narrowing of the midface and midline facial prominence convergence. This phenotypic overlap demonstrates that null mutations of the transition zone predictably drive craniofacial development into a shared, class-wide malformation phenotype precisely as the embryos enter the palatogenesis window. Crucially, this shared structural collapse is specifically tied to the transition zone complex rather than representing a generic consequence of all primary cilia dysfunction. While transition zone complex mutations consistently drive midfacial narrowing and premature convergence [Figure 1B], mutations affecting other ciliary compartments (such as intraflagellar transport (IFT) proteins (e.g., *Ift88* or *Ift140*) or basal body structures) frequently present with distinctly divergent craniofacial dysplasias, including severe midfacial widening (Brugmann *et al*., 2010; Schock & Brugmann, 2017). This morphogenetic contrast firmly establishes that the spatial compression and resulting mechanical trap described here are unique, class-specific signatures of transition zone disruptions.

**Figure 1.**
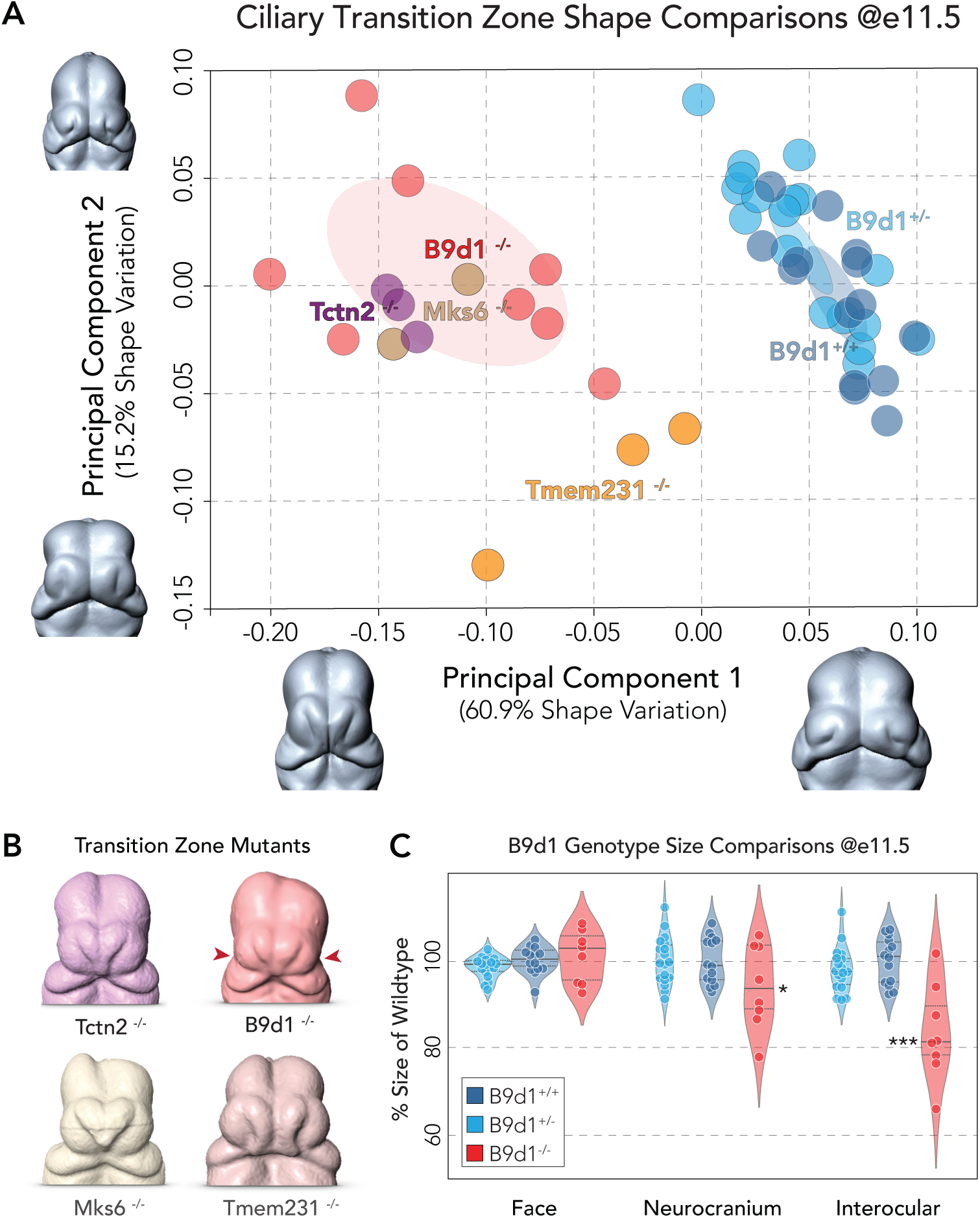
Morphometric analysis of cilia mutant mouse faces. (A) Principal Components Analysis (PCA) of unconstrained external craniofacial shape variance across a multi-mutant cohort at e11.5. Null mutants (*B9d1* [red]*, Tmem231* [orange]*, Mks6* [brown]*, Tctn2* [purple]) are separated from the normal wildtype/heterozygotes (blue) trajectory and cluster into a shared, class-wide malformation phenotype characterized by extreme midfacial convergence. Morphological deformations associated with this convergence suggest a severe narrowing of the forebrain at the level of the eye fields relative to the face (cf. PC1 extremes). (B) Example mutant embryos (color-coded, as above) highlight the midline convergence in transition zone mutants at e11.5. (C) Comparison of centroid size and interocular width across wildtype/heterozygotes and homozygous *B9d1* mutant embryos at e11.5 (violin plots, median (solid line) and interquartile range (dashed lines) are shown). Facial prominence size remains robust in all genotypes, while transition zone nulls exhibit significant mean reduction in brain size (*p<0.05) and particularly interocular width (*p<0.001), suggesting a physically narrowed ventral brain is associated with midfacial convergence in this genotype.

### Brain Size Reduction Drives Midface Narrowing

Critically, the specific shape changes driving this mutant clustering provide a direct mechanical clue to the etiology of their shared facial defects. To assess whether the observed dysmorphology originates solely from localized facial tissue failure or is compounded by broader structural constraints, we compared the absolute sizes (centroid size) of the facial prominence, neurocranium, and interocular width in wild-type (N=14), heterozygous (N=18), and B9d1 (N=8) mutant embryos at this e11.5 fusion window [**Figure 1C**]. This analysis revealed that the absolute size of the facial prominences remains remarkably robust in the mutants relative to normal and heterozygous controls (post-hoc Dunnett’s test, WT vs. Mutant, p=0.641). In contrast, the absolute size of the B9d1 neurocranium is significantly reduced relative to wildtype (post-hoc Dunnett’s test, WT vs. Mutant, p=0.040), particularly at the level of the eye field (post-hoc Dunnett’s test, WT vs. Mutant, p<0.000).

This macro-level spatial loss aligns with recent cellular findings demonstrating that the loss of ciliary Shh signaling in transition zone mutants causes widespread and early apoptosis within the ventral neuroectoderm (Abrams & Reiter, 2021). Our data further reveals the structural consequence of this localized neural cell death: a physically stunted neurocranial scaffold. This is a critical distinction since although Shh signaling is highly active within the facial prominences themselves, the ciliary effect on macro-level tissue growth appears heavily skewed toward the brain module, as evidenced by the lack of change in the size of the facial module compared to normal controls. Thus, while ciliary transition zone mutations have been described as driving a “midfacial” defect, this evidence suggests that the brain instead is the primary physical driver of the observed facial phenotype.

Given that the expanding embryonic forebrain acts as the structural landscape upon which the facial prominences grow, we further hypothesized that a smaller neurocranium acts as a spatial constraint, which would by e12.5 drive even further narrowing and dysmorphology of the transition zone mutant midface. Specifically, we sought to test whether reduced forebrain width physically compresses the morphogenetic field, prematurely closing the spatial gap between growing prominences and forcing atypical premature fusion. To rigorously test this spatial compression hypothesis, we projected the high-resolution internal anatomy of the *B9d1* mutant onto a continuous developmental morphospace of normal murine palatogenesis.

### The Normal Developmental Morphospace and the Fusion Bottleneck

To establish a normal baseline, we next quantified the developmental morphospace of the face and brain from e9.5 to e12.0 in C57bl wildtype mice. PCA of both the facial and neurocranial shape coordinates reveals that the primary axis of variation (PC1) strongly and positively correlates with developmental age, capturing embryonic growth [**Figure 2A-B**]. However, mapping the secondary axis of variation (PC2) against continuous developmental age reveals a non-linear allometric switch [**Figure 2A-B, left axis**]. Both the face and the brain execute a morphospace pivot — a trajectory “U-turn” — that occurs between e11.0 and e11.5, consistent with other amniote species (Young *et al*., 2014; Hu *et al*., 2026). This shape window coincides with the chronological timing of primary palatogenesis and the fusion of the facial prominences in the mouse.

**Figure 2.**
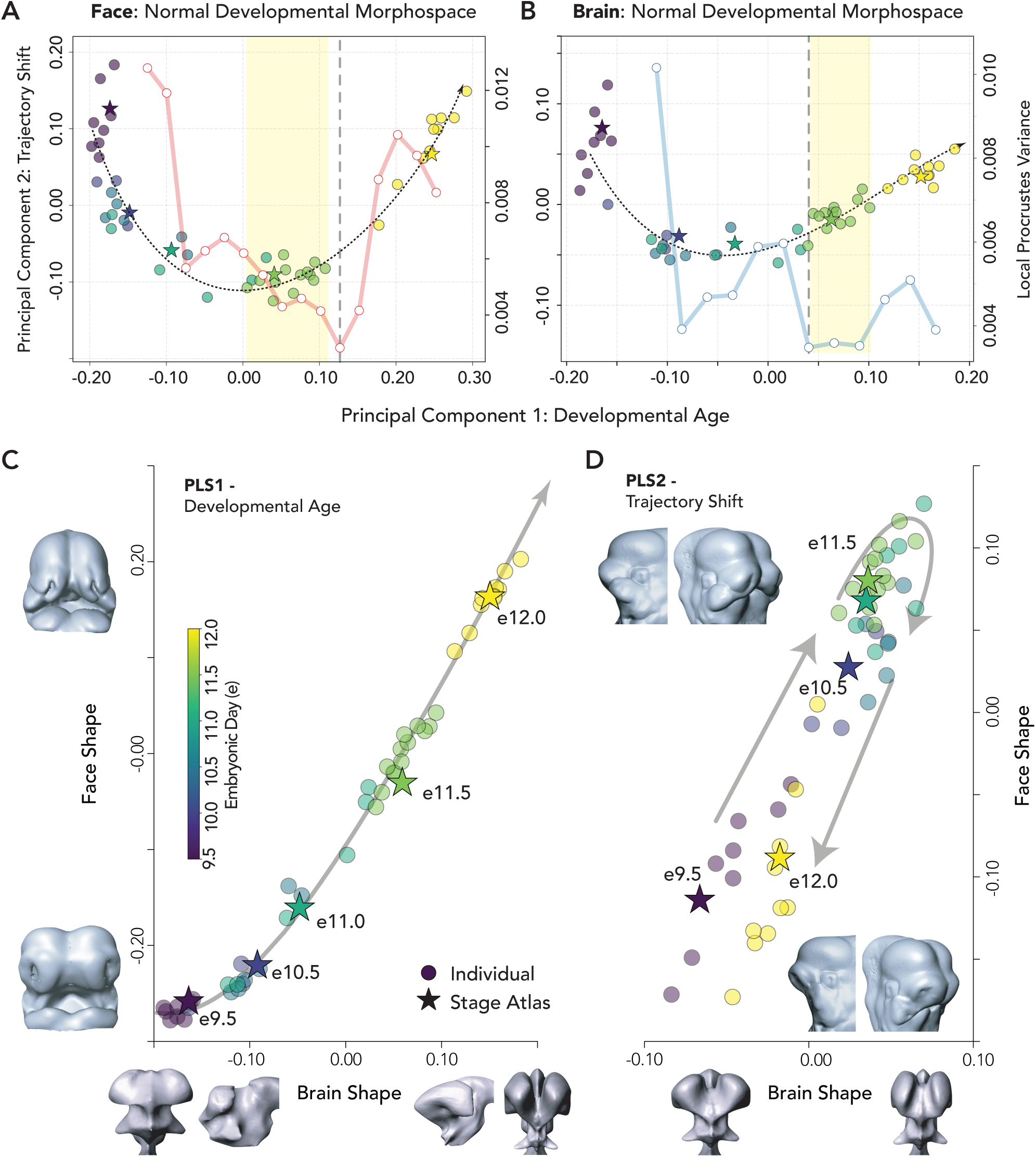
The Normal Mouse Developmental Morphospace, Fusion Bottleneck, and System-Wide Morphological Integration. (A-B) Principal Components Analysis (PC1 vs. PC2) of the normal murine developmental series (e9.5–e12.0) for the Face (left) and Brain (right) modules. In both analyses, PC1 captures developmental age, while PC2 isolates a synchronized, non-linear morphospace trajectory shift (dashed line) occurring ∼e11.0–e11.5 fusion window (yellow box). The continuous sliding-window analysis (ΔPC1 width=0.10, step=0.05) of local Procrustes variance (right axis) shows both the face (solid red line) and brain (solid blue line) exhibit a collapse in morphological variance (minimum=heavy dashed line) around the time of fusion, defining a restrictive “shape window” during palatogenesis. Subsequently, facial variance rebounds as the prominences grow into the mature palate while brain variance remains low due to spatial encapsulation within the neurocranium. (C-D) Two-block Partial Least Squares (PLS) analysis between the developing normal brain and face demonstrates significant spatiotemporal covariation and integration. PLS1 accounts for 97.7% of shared shape covariation (r=0.984), indicating that brain and face growth are tightly coordinated. PLS2 shows that brain expansion and facial convergence reverse their trajectories together at fusion, ∼e11.0-11.5. Axes illustrate extremes of shape variation associated with PC1and PC2.

To test the structural constraints associated with this morphospace trajectory shift, we next applied a sliding-window analysis to track continuous morphological variance (local Procrustes variance). We observed that the facial module displays a classic “hourglass” model of developmental constraint [**Figure 2A-B, right axis**]. As facial prominences grow independently during the early pre-fusion stages (e9.5–e10.0), local variance is elevated. However, by e11.0–e11.5 window, spatial variance descends to a minimum. This system-wide variance collapse supports the presence of a strict structural bottleneck required for successful soft-tissue tethering, after which facial shape variance begins to rebound as the post-fusion face initiates divergent outgrowth. Variance in the brain similarly collapses into facial fusion, but thereafter it fails to increase further, likely due to spatial encapsulation rather than outgrowth (Hu *et al*., 2026).

### System-Wide Morphological Integration

To verify that the simultaneous variance collapses and morphospace pivots in the face and brain were not coincidental, we assessed their spatiotemporal covariation using a two-block Partial Least Squares (PLS) analysis from e9.0-12.0 [**Figure 2C-D**]. The normal developmental series exhibits exceptionally high integration between the two tissue modules. The primary axis of covariation (PLS1) accounts for most of the shared shape variation, demonstrating a near-perfect correlation (r = 0.984) between the brain and the face. This intense covariation supports the hypothesis that the structural expansion of the forebrain and the convergence of the facial prominences operate as a tightly coupled and highly synchronized morphogenetic system.

### Transition Zone Complex Mutations Drive Heterochronic Uncoupling

Having established the integrated baseline of normal development, we tested our hypothesis that transition zone complex mutations (B9d1) disrupt the spatiotemporal coordination required for typical fusion. Using a mathematical projection approach, we transformed the shape coordinates of e11.5 B9d1 mutant embryos into the PCA space defined by the normal mouse ontogenetic shape trajectory [**Figure 3A-B**]. The projection revealed a heterochronic uncoupling between the two tissue modules. In the brain morphospace, the normal e11.5 mean PC1 score is ∼-0.07, but the B9d1 mutants exhibit a significantly lower mean score (∼0.01; Student’s t-test, p<0.001) [**Figure 3A**]. This leftward shift along the developmental axis indicates a decelerated brain shape that morphologically resembles a less mature embryo (∼e11.0) [**Figure 3C**]. Conversely, in the facial morphospace, the normal e11.5 mean PC1 score is -0.07, while the B9d1 mutants exhibit an increased mean score (0.05; Student’s t-test, p<0.001) approaching the normal e12.0 mouse (0.13) **[Figure 3B].** This rightward shift indicates an accelerated facial geometry, wherein the normally sized prominences have been mechanically forced into a prematurely convergent spatial configuration typical of older embryos [Figure 3D]. Furthermore, supporting the prediction that mutants retain structural correlation but suffer from destabilized spatial constraints, the shape variance for the mutant cohort was significantly higher than that of the wild-type cohort at e11.5 for both the brain and face (Levene’s test, p<0.001).

**Figure 3.**
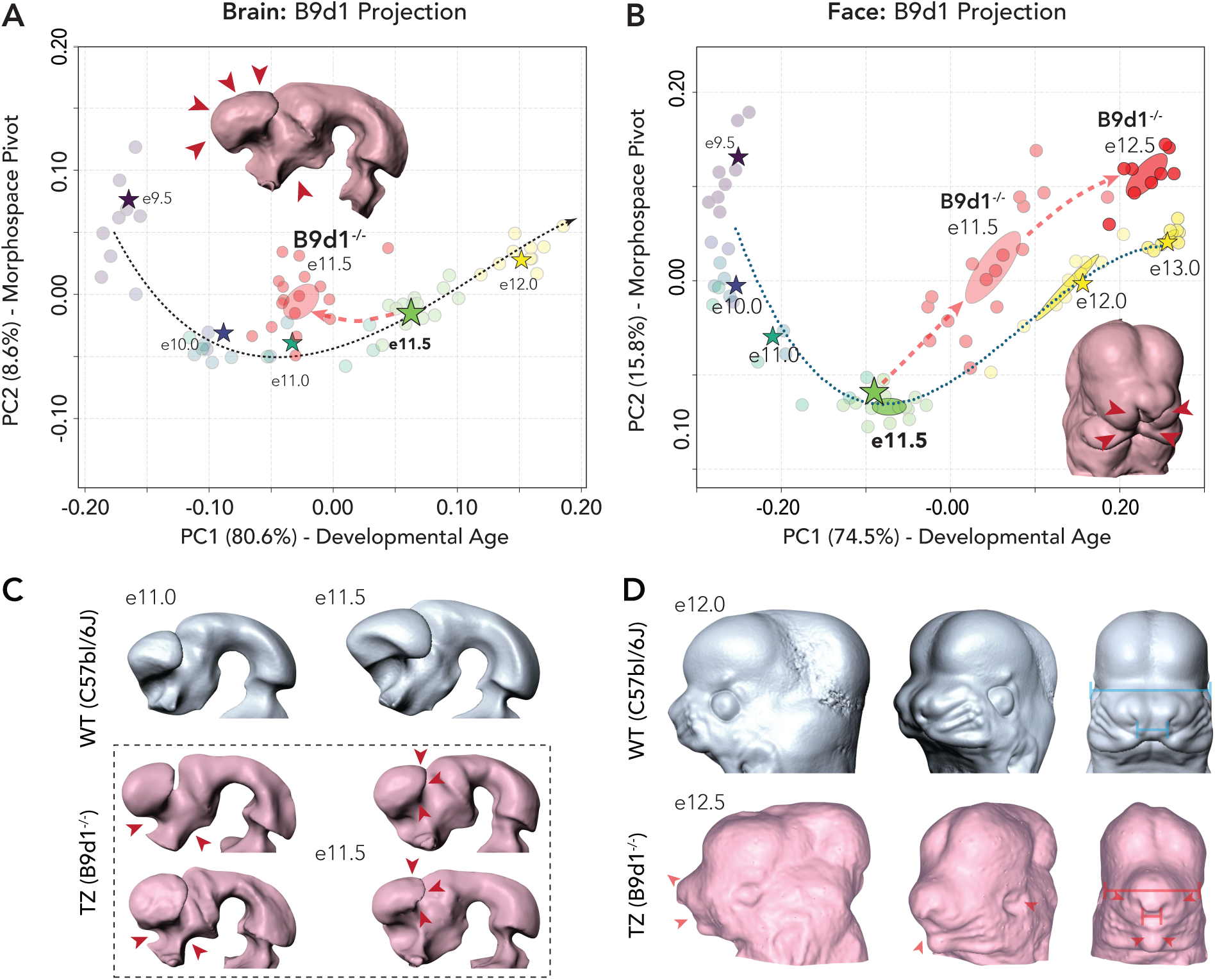
Heterochronic Uncoupling in the Mutant Phenotype. (A-B) Projection of *B9d1* mutant embryos (light red = e11.5; dark red = e12.5) into the normal developmental morphospace demonstrates a heterochronic mismatch. Brain shape at e11.5 is decelerated (shifted left, less mature), while facial shape is accelerated (shifted right, more mature) relative to age-matched wildtype, mechanically forcing intrinsically normal prominence growth into premature apposition, a phenotype that persists into e12.5 mutants (dark red). (C). Representative examples of normal wildtype brain morphology at e11.0 and e11.5 (top row, blue) contrasted with those of B9d1 mutant morphology at e11.5. Mutants variably exhibit reduced ventral diencephalic structures posterior to the neurohypophyseal bud and anterior to the optic chiasm (left, red arrows), but consistently exhibit reduced telencephalic cerebral vesicles more consistent in shape and size with e11.0 wildtypes (right, and all specimens). (D). When compared to normal e12.0 mice (blue, top), e12.5 mutant faces (red, bottom) retain a narrower midface driven by the premature convergent distal apposition and fusion of the maxillaries (lines = illustrate representative widths), conditioned on slower forebrain growth.

To definitively assess the persistence and physical nature of this heterochronic shift, we further evaluated the faces of an older cohort of *B9d1 m*utants at e12.5 (N=9). Projecting these mutants into the normal morphospace revealed that they advance even further along the primary axis, maintaining the highly fused, mature geometric configuration. Crucially, despite this structural convergence, the intrinsic size of the e12.5 mutant facial prominences remains statistically identical to e12.5 wildtype controls (post-hoc Dunnett’s test, wildtype versus mutant, p=0.368). This confirms that the premature fusion is a secondary spatial consequence of the collapsed brain scaffold, uncompromised by intrinsic tissue loss.

## Discussion

### A System-Wide Shape Window for Palatogenesis

Our geometric morphometric characterization of normal mouse craniofacial development from e9.5–e12.0 establishes a rigid system-wide morphogenetic constraint dictating the physical environment of primary palatal fusion. The sliding-window variance analysis reveals an “hourglass” bottleneck, where morphological variance within both the face and the brain collapses between e11.0 and e11.5. This drop in spatial variance aligns with the timeline of facial prominence apposition and fusion, mirroring our comparative findings in avians (Hu *et al*., 2026).

The integrated brain-face developmental morphospace confirms that successful palatogenesis depends on the entire craniofacial system navigating a narrow highly constrained “shape window.” Prior to e11.0, the facial prominences grow independently and possess substantial morphological flexibility. Upon contact and fusion (e11.0-11.5) that structural freedom collapses. However, this shape window is not merely defined by the independent actions of the facial prominences but rather two-block PLS analysis indicates an intense system-wide integration that represents a highly synchronized physical event coordinated with brain growth. This covariation supports the hypothesis that the structural expansion of the forebrain acts as the physical conditional landscape in which facial contact and fusion occur, defining the temporal and spatial bounds of normal and abnormal palatogenesis. This intense macro-level covariation provides a a modern 3D quantitative counterpoint framework for the foundational concept originally proposed by DeMyer and subsequently expanded by Muenke that “the face predicts the brain” (DeMyer *et al*., 1964; Muenke *et al*., 1994).

### Heterochronic Uncoupling and Premature Fusion in B9d1 Mutants

Projecting the B9d1 (**transition zone** complex) mutant embryos into the normal PCA morphospace provided a testable mechanism to determine how disruptions to a constrained shape window induce craniofacial defects. Our results indicate that the B9d1 mutation does not cause a generalized global developmental delay or simple reductions to absolute prominence size, rather it drives a heterochronic uncoupling between the integrated brain and face modules, and it is this disruption that imposes a severe spatial constraint that strongly correlates with downstream dysmorphology.

At e11.5, the B9d1 null brain is significantly smaller and narrower than normal controls while the facial prominences remain comparable in size to the wildtype. Consequently, mutant embryos are shifted in both brain and face developmental morphospaces, reflecting an immature brain and abnormal fusion of the maxillaries in the midline. This heterochronic shift explains the mechanical origin of the premature fusion events characteristic of this mutation. Because the underlying brain landscape is physically stunted and structurally narrowed, the physical distances required for contact of the prominences are likewise reduced. At the same time, the growth and size of the facial prominences is less affected, so they continue to develop normally but in an unusually reduced landscape, ultimately leading to premature apposition and atypical fusion. Critically, this mechanistic system-wide perspective provides a system-wide framework for previously proposed cellular models of ciliary transition zone defects. Recent work characterizing midfacial defects in transition zone mutations (including Tctn2 and Mks6) has demonstrated that premature fusion involves early apoptosis within the ventral neuroectoderm as well as localized cell death within frontonasal mesenchyme derived from the cranial neural crest (Abrams & Reiter, 2021). According to this cellular model, the loss of ciliary mediated SHH-signaling increases apoptosis in the midline leading to a midline collapse.

Our 3D quantitative analysis complements this finding by separating the disparate morphogenetic consequences of this apoptosis. Despite localized cell death in the midline, the centroid size of the B9d1 mutant facial prominences at the critical e11.5 fusion window remains remarkably robust compared with wildtype. In contrast, the spatial loss is mainly confined to the forebrain. Placing the altered facial shape within a continuum of morphological space, our data strongly suggest that midfacial narrowing is driven by a synergistic, multiscale mechanism. The strongly localized midline deficits occur in parallel with a facial complex that develops on an abnormally narrowed neurocranial scaffold. The smaller forebrain, which is itself a result of early enhanced neuroectodermal cell death, essentially acts as a mechanical trap, exacerbating the collapse of the midline. This macro-level spatial constraint aligns with our previous findings demonstrating that physical interactions and tissue-tissue spatial proximities between the brain and face are as critical to normal morphogenesis as their shared molecular signaling pathways (Marcucio *et al*., 2011; Chong *et al*., 2012). While our quantitative spatial data strongly supports this model, future studies utilizing conditional knockouts to genetically separate the neural and neural crest lineages will be necessary to definitively isolate this physical mechanism from complex cross-tissue molecular signaling.

Furthermore, this modular disparity highlights the complex spatiotemporal dynamics of SHH signaling across the developing head. While SHH signaling is essential for the development of both the viscerocranium (facial prominences) and the neurocranium (forebrain), by e11.5 the structural consequences of transition zone complex mutations are unevenly distributed and strikingly specific. Disruption of ciliary gating causes sufficient ventral neuroectodermal apoptosis to inhibit brain shape, yet the adjacent facial processes maintain normal growth and volume. This suggests that the requirements for SHH signaling induced by ciliary mutation vary significantly in time and space between adjacent tissues. At the onset of palatogenesis, the brain and face display divergent macro-level vulnerabilities to the transition zone mutation, suggesting that facial mesenchyme may possess distinct volumetric signaling thresholds, rely on alternative pathway redundancies, or operate on a delayed macro-level timescale of ciliary dependence compared with the neighboring neural ectoderm. Consequently, what clinically presents as a single “midfacial” abnormality is instead caused by localized, module-specific signaling disruptions that cascade into a system-wide physical mismatch.

### The Continuum of Premature Fusion and Cleft Lip

These findings offer a novel unified framework for conceptualizing craniofacial dysmorphologies such as premature fusion and non-syndromic cleft lip [**Figure 4A-C**]. Rather than viewing these defects solely as localized failures within the facial prominences themselves, our data strongly suggest that instead contact and fusion event outcomes reside at extremes of a continuous phenotypic spectrum conditioned by brain development (Boughner *et al*., 2008; Chong *et al*., 2012).

**Figure 4.**
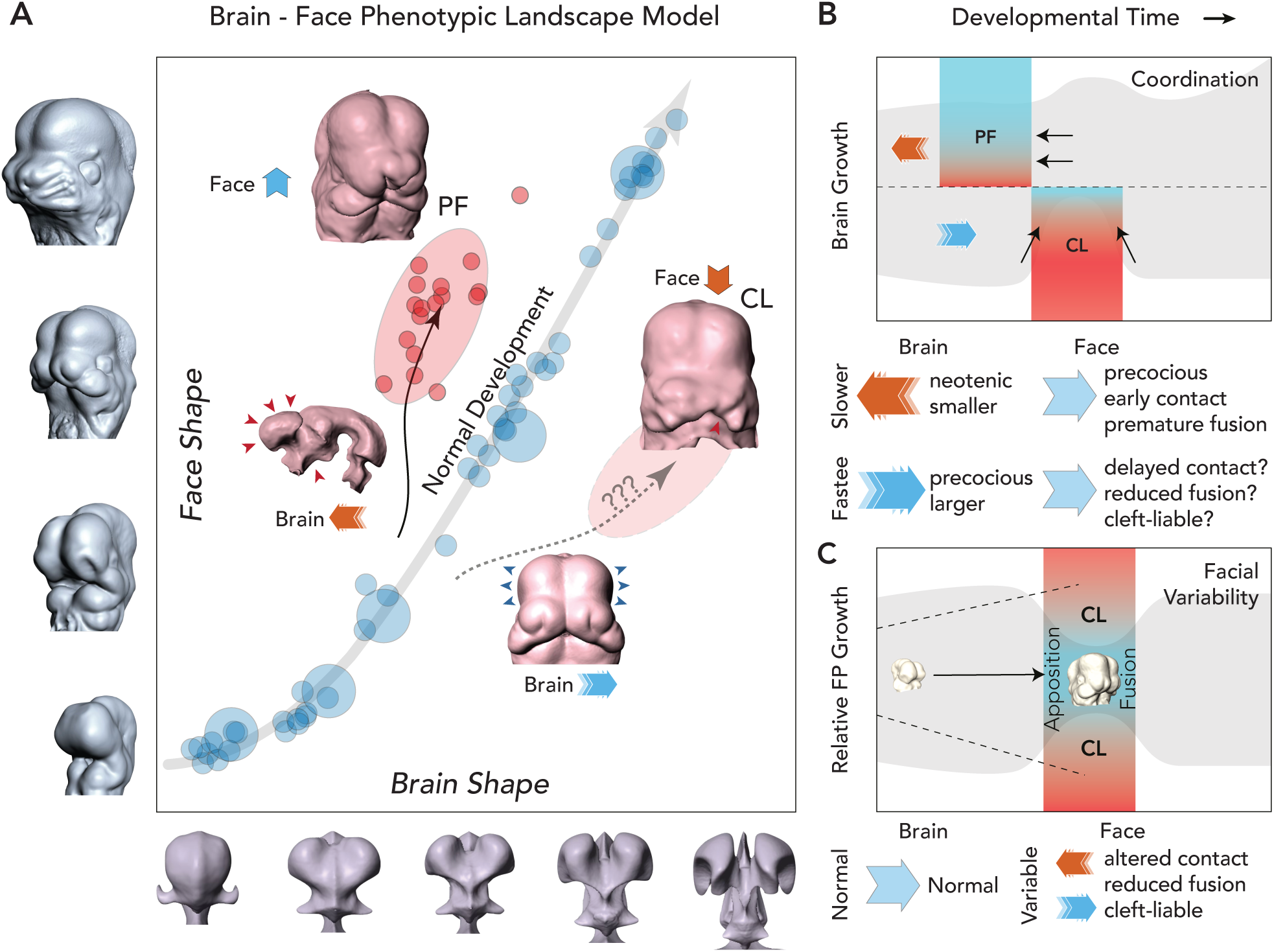
A Unified model of the Brain as a Conditional Phenotypic Landscape. A. Brain shape and size impacts the physical distance that facial prominences must cross to achieve successful apposition, merging, and fusion with each other, a process that is highly coordinated in normal development (shown, PLS of the wildtype developmental morphospace including B9d1 mutants). If brain growth is decelerated this threshold is reduced, in which case normal prominence growth results in Premature Fusion (PF) in the midline, as found in *B9d1* mutants [red]. Conversely, if brain growth is accelerated, we hypothesize that the expanding forebrain will act as a physical wedge, creating a spatial gap that increases the likelihood of Cleft Lip (CL) outcomes. B. Schematic representation of the brain-face predictions showing how acceleration and deceleration of brain growth alter the predicted landscape of apposition and fusion (grey shading), either expanding or reducing this region of morphospace. (C) When brain growth is normal successful facial prominence apposition and fusion is dependent on navigating a narrow shape window, thus elevated morphological variance or altered relative growth ratios between the prominences may push the tails of the population distribution outside the required bounds for fusion, increasing the likelihood of cleft lip (Young *et al*., 2014, Hu *et al*., 2026).

During normal brain development, the most critical factors determining contact and fusion are dependent on the facial prominences themselves. Our model suggests that successfully navigating the spatial “shape window” requires three intact physical parameters: scaffold, vector, and volume. When the underlying brain scaffold is abnormally slow, narrow, or restricted (as in the B9d1 mutant), the facial prominences will span the diminished landscape too rapidly, resulting in premature fusion. Conversely, if the brain scaffold expands faster than normal, it acts as a physical wedge, increasing the physical distance required to achieve contact and increasing the likelihood of cleft lip (Boughner *et al*., 2008; Marcucio *et al*., 2015). This mechanical principle is empirically supported by *Ift88* ciliary mutants, where severe midfacial widening physically prevents midline prominence convergence, directly resulting in clefting (Brugmann *et al*., 2010). Even with a normal scaffold, fusion can fail if the geometric vector is misdirected (as seen in the *Unicorn* mutant, where prominences physically bypass the fusion window) (Lantz *et al*., 2020), or if the prominences lack the cellular volume to bridge the gap (as seen in the *Srsf3* mutant, where severe apoptosis stalls outgrowth) (Dennison *et al*., 2021). Future efforts to test this corollary are ultimately required to establish the model’s ability to predict a range of outcomes.

In conclusion, our continuous DM approach highlights how developmental constraints dictate the structural boundaries of palatogenesis. By decoupling the morphogenetic trajectories of the brain and face, TZ-complex mutations explicitly highlight that the size, shape, and timing of the underlying neurocranium are dependencies for successful facial fusion, offering a multi-scale mechanical explanation for the spectrum of clinically observed midfacial defects.

### Study limitations

Several limitations should be considered when interpreting these findings. First, while our quantitative morphometric projection strongly supports a mechanical etiology for premature fusion, our reliance on global transition zone null mutant models (*B9d1^-/-^*) restricts our ability to definitively prove tissue-autonomous causality. Differentiating the pure physical spatial constraints imposed by the stunted neurocranium from complex, secondary cross-tissue molecular signaling deficiencies will ultimately require tissue-specific conditional knockouts (e.g., neural crest versus neuroectodermal deletions). Second, the retrospective nature of our high-resolution micro-CT cohort precluded the collection of precise physiological staging metrics, such as tail somite counts. To mitigate intra-litter chronological variation, we utilized the primary axis of shape variance (PC1) as an objective proxy for developmental age. While this is a highly validated and robust standard in continuous 3D geometric morphometrics, future models utilizing exact somite-matched pairings could provide an even finer temporal resolution of the fusion bottleneck. Finally, our characterization of heterochronic uncoupling defines facial “advancement” strictly through macro-level spatial geometry and premature structural convergence. Because this architectural analysis was not paired with parallel spatial transcriptomics, we cannot completely rule out the presence of subtle, localized alterations to the intrinsic gene regulatory networks within the facial prominences. Integrating these continuous 3D shape trajectories with targeted molecular tracking will be essential to fully map the intersection of physical and cellular dynamics driving palatogenesis.

## Materials and Methods

### Animal Husbandry and Embryo Collection

All animal experiments were conducted in accordance with approved Institutional Animal Care and Use Committee (IACUC) protocols (UCSF, AN207759-00B). transition zone mutants (*B9d1*, *Tmem231*, *Mks6*, and *Tctn2*) were maintained on a standard C57bl background. Timed matings of heterozygous (^+/-^) mice were utilized to generate both normal (wild-type and heterozygous) and homozygous mutant (^-/-^) embryos. Noon on the day a vaginal plug was observed was designated as embryonic day 0.5 (e0.5). To establish a continuous, high-resolution baseline of normal murine palatogenesis, normal embryos (C57bl/J) were collected at half-day intervals from e9.5 through e13.0. Because gestational plug dates are susceptible to intra-litter temporal variation, the primary axis of shape variance (PC1) was utilized throughout the analysis as the quantitative proxy for true morphological developmental age (Musy *et a*l., 2018; Hu *et al*., 2026). Homozygous transition zone mutant embryos were specifically harvested at e11.5, the critical chronological window for primary palatal fusion in the mouse, and at e12.5 to assess the persistence of the facial phenotype. Following harvest, embryonic yolk sacs were collected for PCR genotyping. The initial sample of transition zone embryos were fixed in 4% paraformaldehyde (PFA) in PBS, while the normal mouse sample and subsequent B9d1 samples were additionally stained in a 1% Lugol’s iodine contrast solution to enhance the radiopacity of embryonic soft tissues for downstream imaging (Degenhardt *et al*., 2010; Wong *et al*., 2013).

### Micro-Computed Tomography Imaging, Registration, and Segmentation

Embryos were imaged via high-resolution micro-computed tomography (μCT, Scanco VivaCT 40) at a resolution of 10μm. To establish baseline morphological targets, we utilized an automated, non-linear volume-based registration pipeline to generate stage-specific mean volumes (Percival *et al*., 2019). Source images were spatially aligned to an iteratively defined average target image using hierarchical affine and non-linear deformations via the ANIMAL algorithm. For every individual sample and stage-specific consensus volume, we generated two highly detailed 3D segmentations: (1) an external surface isolating the developing “face” (facial prominences and oral cavity) and (2) an internal volume isolating the “brain” by approximating the dense, underlying neuroepithelium [**Supplemental Figure S1**].

### Landmarking & Geometric Morphometrics (3DGM)

A standard, reproducible set of 3D anatomical landmarks similarly apportioned across modules was placed on the stage-specific mean volumes of the face (N=40) and brain (N=43), serving as a unified morphological atlas [**Supplemental Figure S2**]. A semi-automated landmark estimation protocol (*Landmark Editor* v.3.01, UC Davis) was then applied to rapidly identify homologous points across all individual normal and mutant embryos, followed by rigorous manual adjustment and verification (Devine *et al*., 2020). All landmarks were manually verified and adjusted as needed by a single observer. To definitively assess landmarking reproducibility, intra-observer error was calculated via Procrustes ANOVA on a subset of repeated specimens, confirming that measurement error was negligible (<1%) relative to total biological shape variation. The raw 3D coordinate configurations were subjected to Generalized Procrustes Analysis (GPA) to mathematically quantify shape variation separate from the confounding effects of scale, position, and spatial orientation (Zelditch *et al*., 2004; Adams & Otarola-Castillo, 2013).

### Morphospace Construction and Continuous Shape Variance

Within-species developmental morphospaces (DMs) for both the face and brain modules were quantified using Principal Components Analysis (PCA) of the Procrustes shape coordinates (Mitteroecker & Gunz, 2010). To assess structural constraints, we quantified continuous morphological variance across the normal developmental series using a sliding-window analysis (Hu *et al*., 2026). Normal specimens were sorted by PC1, serving as an objective proxy for continuous developmental age. A sliding window of fixed PC1 width (ΔPC=0.10) was translated along the PC1 axis (step=0.01), dynamically calculating a local consensus shape at each step. True morphological variance (local Procrustes variance) was computed as the sum of squared Procrustes distances of the specimens to their localized consensus, yielding a continuous measure of morphospace constraint orthogonal to the primary growth trajectory.

### Morphological Integration (Partial Least Squares)

To test for spatiotemporal covariation and system-wide integration between the developing face and brain, we utilized a two-block Partial Least Squares (PLS) analysis (Rohlf & Corti, 2000; Klingenberg, 2009). The jointly superimposed Procrustes coordinates of the normal developmental series (e9.5–e12.0) were separated into two structural blocks, with block 1 assigned to the brain and block 2 assigned to the face, to extract the primary axes of morphological covariation. This analysis allowed us to quantify the degree of correlation between the two tissue modules across normal developmental time and provided a baseline framework to evaluate whether the B9d1 mutants deviate from this tightly integrated trajectory.

### Mutant Projection and Heterochronic Analysis

To test our hypothesis that TZ-complex mutations induce spatiotemporal uncoupling between the brain and face, we utilized a mathematical projection approach. First, a baseline PCA was fitted strictly using the Procrustes coordinates of the normal developmental series (e9.5–e12.0 in the brain, e9.5-e13.0 in the face) to define the standard ontogenetic trajectory and morphospace boundaries. Next, the Procrustes shape coordinates of the B9d1 mutant embryos (chronological age e11.5-e12.5) were mathematically transformed (projected) into this established normal morphospace. The primary axis of variation (PC1) was aligned to correlate positively with chronological age. We then calculated the mean PC1 scores of the B9d1 face and brain modules and directly compared them against the mean PC1 scores of the normal e11.5 and e12.5 cohort. If present, this projection allowed us to quantify relative heterochronic shifts (i.e., to identify morphological acceleration or deceleration) in the shape progression of the face and brain resulting from the B9d1 mutation.

## Acknowledgements

We thank Jeremy Reiter for sharing primary cilia mutant mouse lines.

## Competing interests

The authors declare no competing interests.

## Funding

This research was funded in part by grants from the National Institutes of Health and the National Institute of Dental and Craniofacial Research: F32DE018596 (NY), R56DE029124 (NY, RM, BH), R01DE018234 (RM, BH), and R01DE019638 (RM, BH).

## Data and resource availability

All data are available upon request.

## Supplemental Information

**Supplemental Figure S1:**
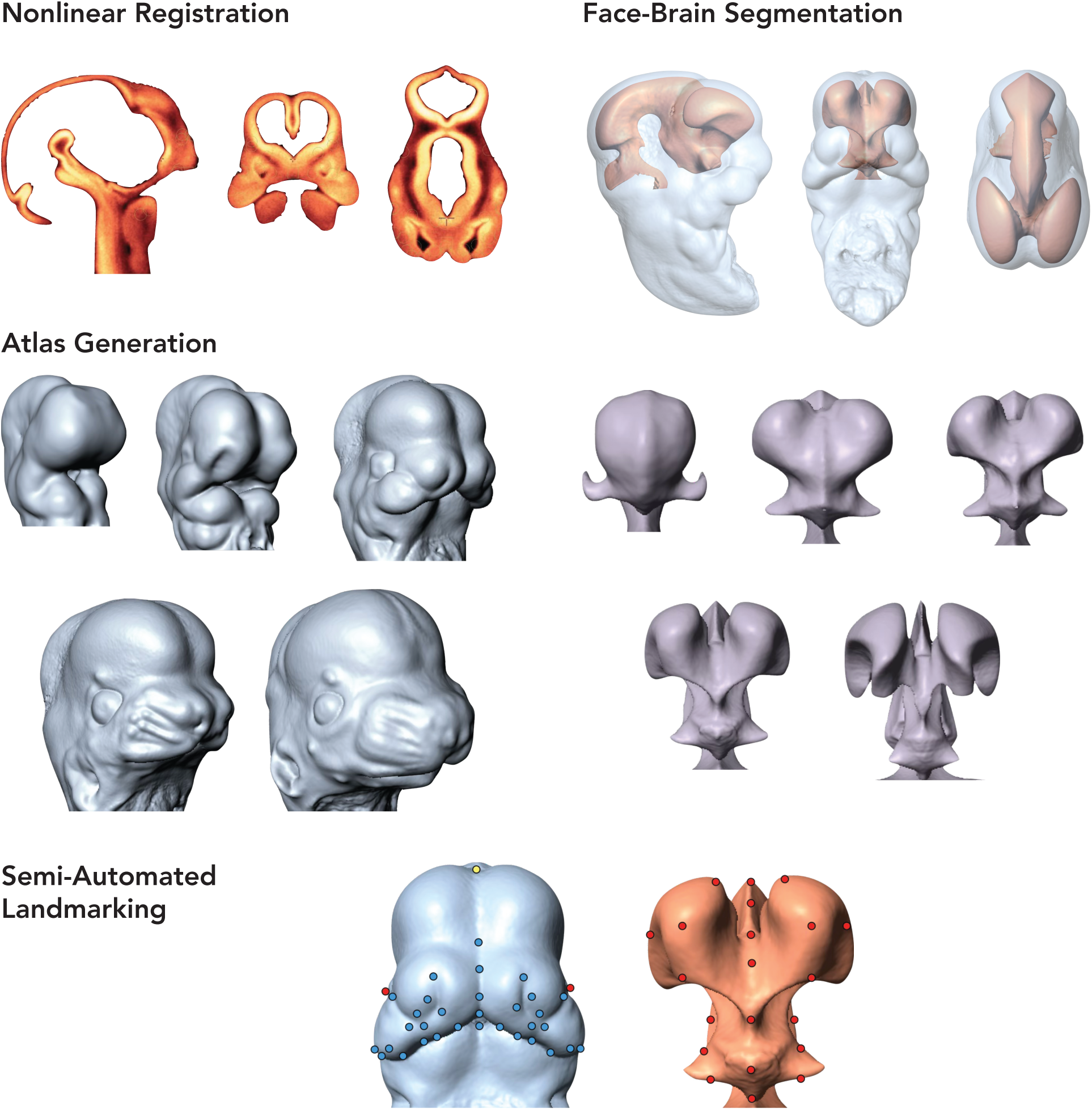
Imaging, Segmentation, and Landmarking Protocol. Representative high-resolution μCT scans of mouse embryos illustrating the workflow from image capture, to nonlinear registration and averaging, segmentation of the external facial ectoderm and internal neuroectoderm, atlas generation, and semi-automated 3D landmarking.

**Supplemental Figure S2:**
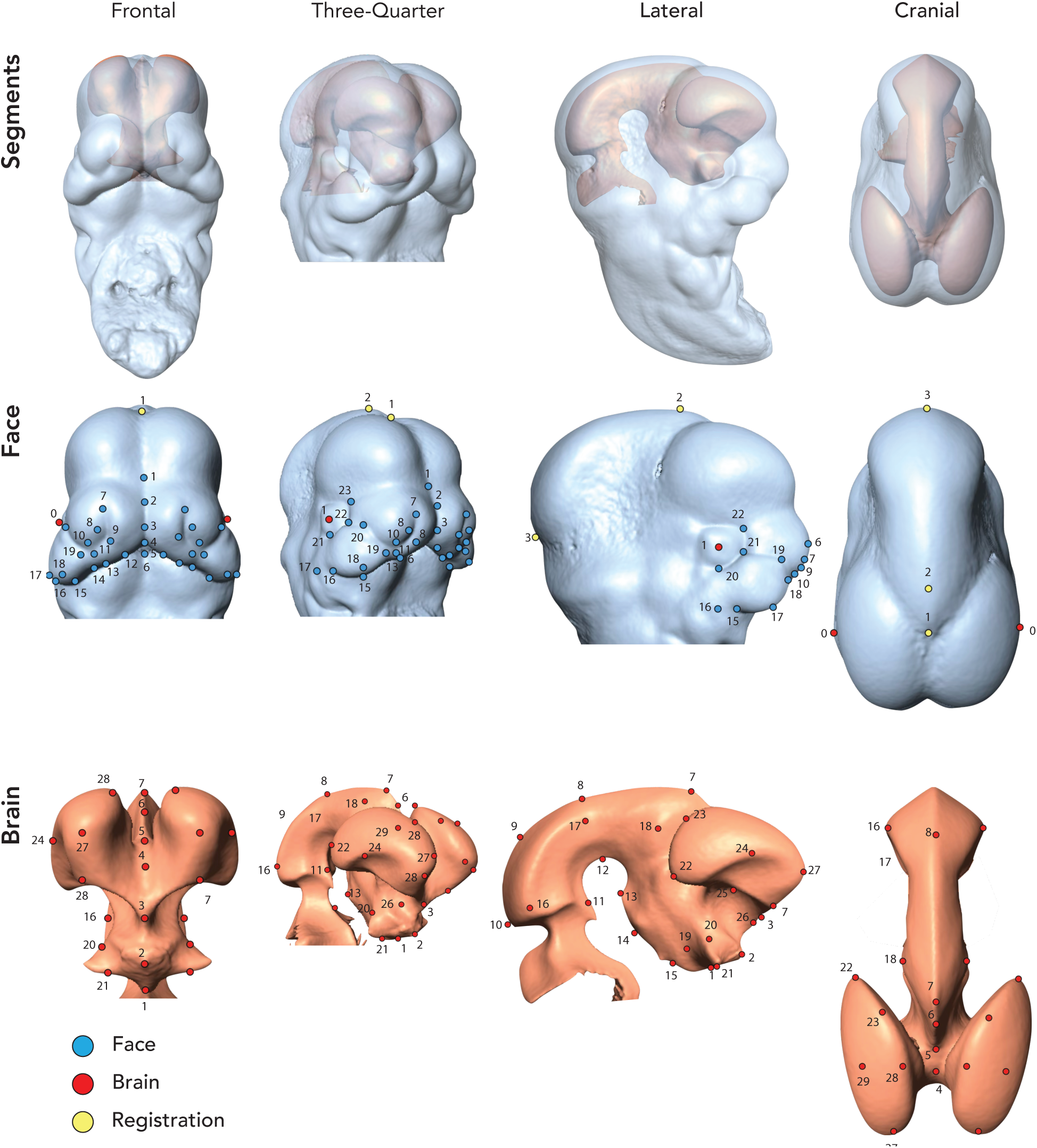
3D Landmarks. Representative high-resolution μCT atlas (e11.5) illustrating segmentations (top) of the external face (second row) and internal neuroepithelium (third row) in various orientations. 3D landmarks are shown for the face (blue) and brain (red) used for Geometric Morphometrics (3DGM), and those used to enhance registration only (yellow). A full description of landmarks can be found in **Supplemental Table S1**.

**Supplemental Table S1:**
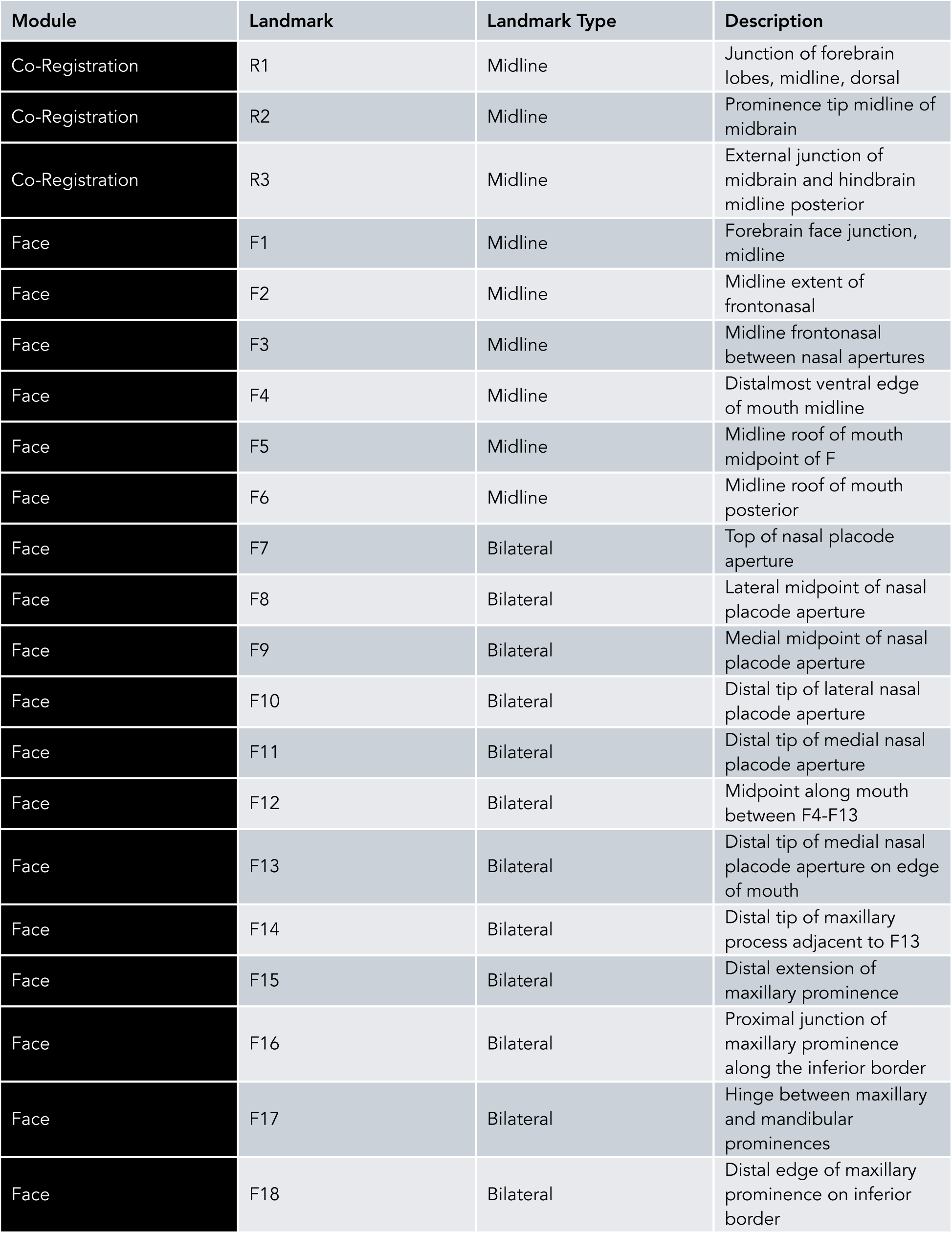
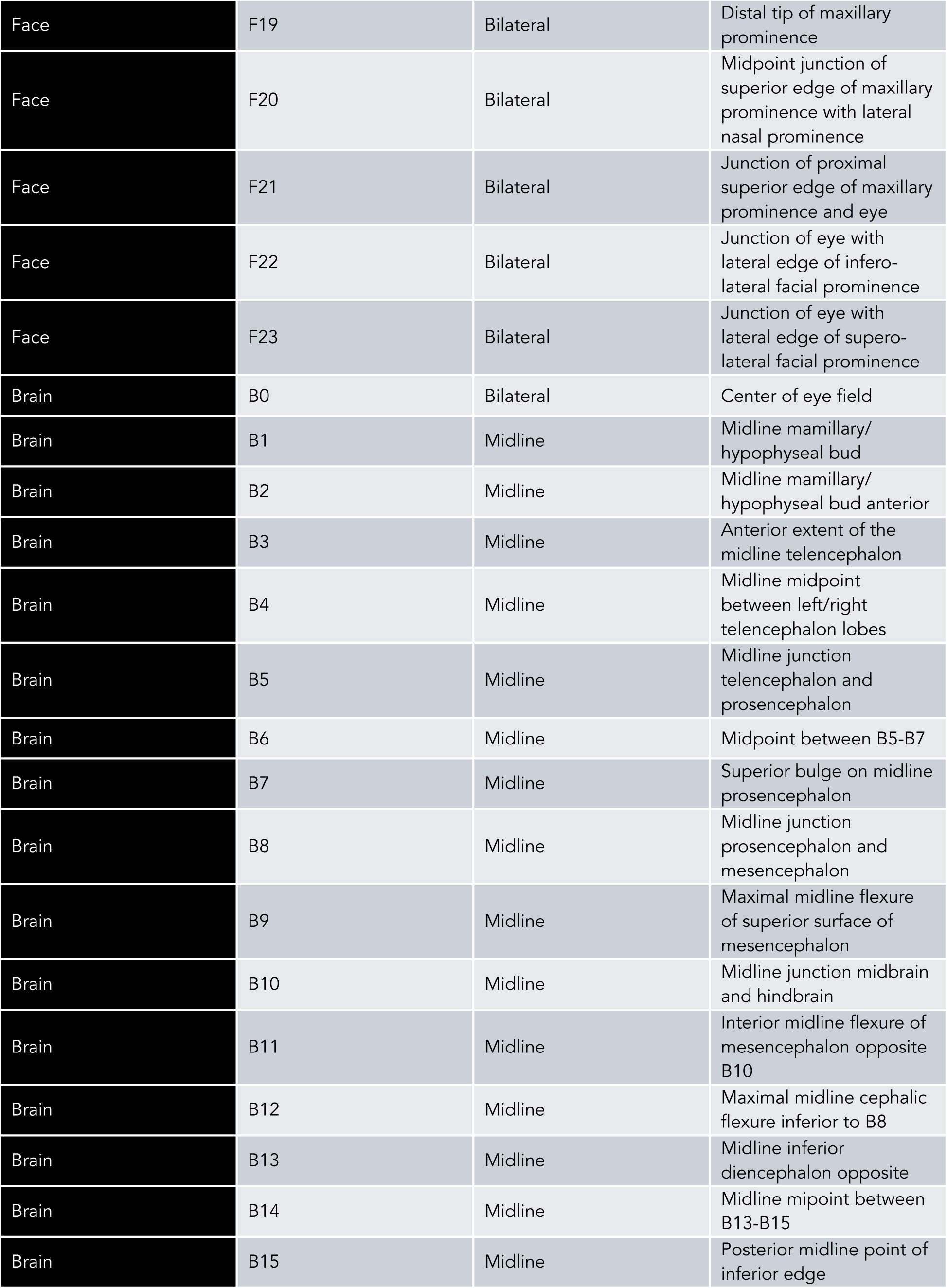
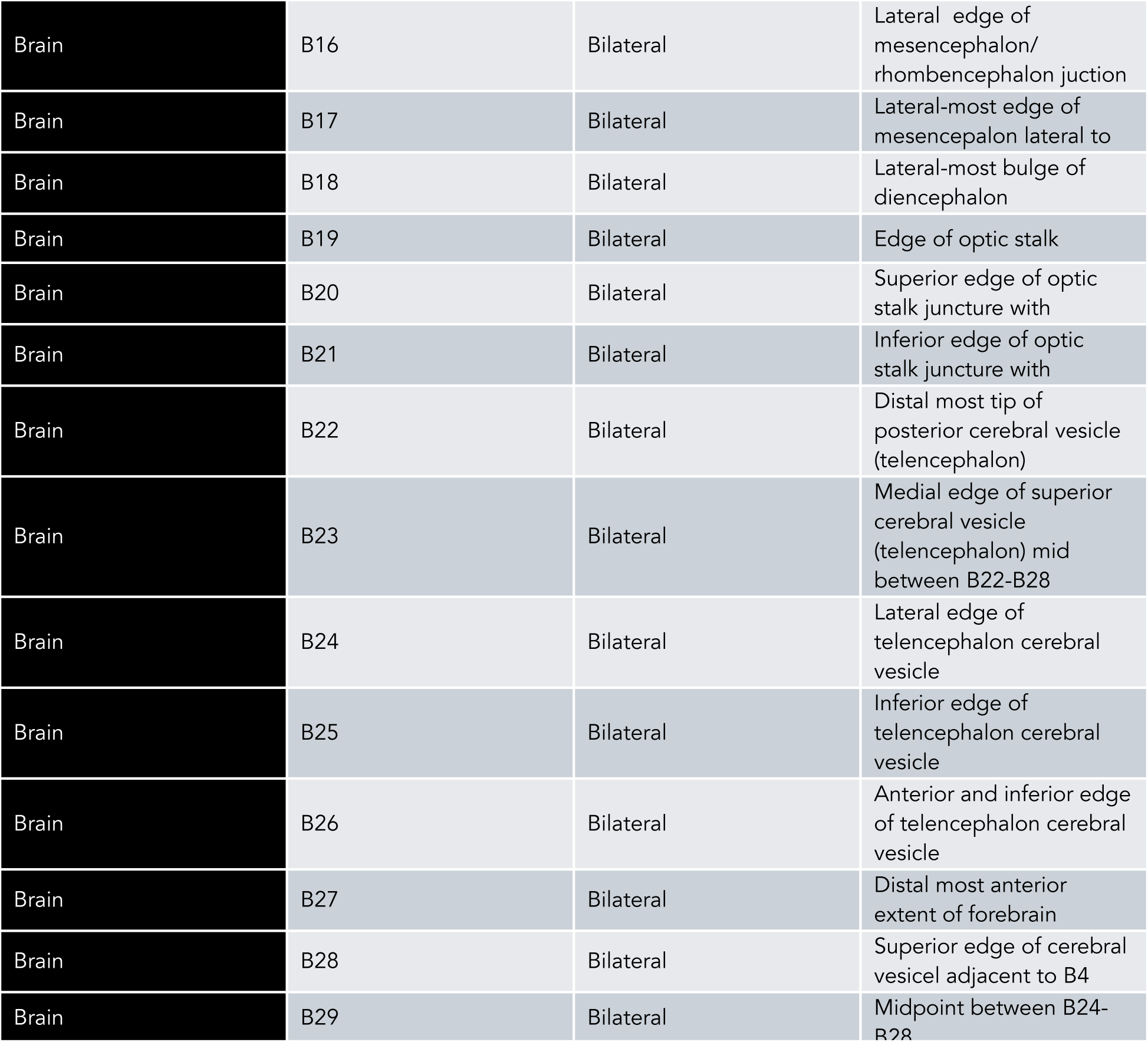
Landmark Descriptions.

## Notes

### Competing Interest Statement

The authors have declared no competing interest.

